# Anomalous diffusion of *E. coli* under microfluidic confinement and chemical gradient

**DOI:** 10.1101/2022.12.12.520016

**Authors:** Md Ramiz Raza, Jijo Easo George, Savita Kumari, Mithun K Mitra, Debjani Paul

**Affiliations:** Department of Biosciences and Bioengineering, Indian Institute of Technology Bombay, Powai, Mumbai, 400076, India; Department of Physics, Indian Institute of Technology Bombay, Powai, Mumbai, 400076, India

## Abstract

We report a two-layer microfluidic device to study the combined effect of confinement and chemical gradient on the motility of wild type *E. coli*. We track individual *E. coli* in 50*μm* and 10*μm* wide channels, with a channel height of 2.5*μm*, to generate quasi-2D conditions. We find that contrary to expectations, bacterial trajectories are super-diffusive even in absence of a chemical (glucose) gradient. The superdiffusive behaviour becomes more pronounced on introduction of a chemical gradient or on strengthening the lateral confinement. Runlength distributions for weak confinement in absence of chemical gradients follow an exponential distribution. Both confinement and chemoattraction induce deviations from this behaviour, with the runlength distributions approaching a power-law form under these conditions. Both confinement and chemoattraction suppress large angle tumbles as well. Our results suggest that wild-type *E. coli* modulates both its runs and tumbles in a similar manner under physical confinement and chemical gradient.

## INTRODUCTION

Bacterial cell behavior generally includes communication, motility and growth. This behavior of a bacterial cell is dependent on its habitat and external stimuli. A hallmark of bacterial habitats is confinement – where bacterial cells exist within restricted boundaries. For example, microcavities in teeth, soil micropores, water filter pores, and endosymbiotic hosts are some well known example of confined spaces[1][2][3][4]. Gut microbiomes are also functionally associated with narrow spaces and chemosensory signaling [5]. The movement of motile bacteria is also influenced by stimuli like chemical gradients, pH, temperature, etc. For example, flagellated *Escherichia coli* (*E. coli*) exhibit “run” and “tumble” patterns in a bulk liquid medium [6]. These motility patterns are affected by chemical cues[7][8][9]. Studying bacteria in controlled environments, where both confinement and chemical gradient can be tuned externally, potentially allows insight into fundamental characteristics of bacteria and how they respond to various physical and chemical cues.

Microfluidic devices are powerful tools to mimic the natural confined habitats of bacteria under controlled experimental conditions [10, 11]. Experiments on *E. coli* in microfluidic devices have yielded much insight into their swimming behaviour. Biondi *et al* [12] characterized *E. coli* motility in 2 *μ*m to 20 *μ*m high channels, and reported an exponential distribution of run times in a 10 *μ*m wide channel. Both runs and tumbles were observed in the trajectories of *E. coli* inside channels of 2 *μ*m width or narrower, with an average velocity of 10 *μ*m/sec across all channels with dimensions larger than the width of a single bacterium [1]. Intermittent hopping and trapping of *E. coli* has been reported for bacteria confined in 3D porous media, with the hop lengths decreasing with decreasing pore size [13]. Tokarova and others [14] explored the motility of different flagellated bacteria including *E. coli* inside microfluidic confinements and concluded that the steric interactions between the flagella and the channel wall drive the motility inside confinements. Interestingly, for high degrees of confinement, it has also been reported that *E. coli* undergo a transition from a *wall-crawling* behaviour where they swim along channel walls, to a more axial swimming mode, where they swim centrally for extreme confinements comparable to *E. coli* dimensions [15].

Bacterial run and tumble motion in free aqueous environments had classically been believed to be diffusive over long timescales [7][9]. However, experimental studies of counterclockwise (CCW) vs. clockwise (CW) rotations of a single motor for an immobilised bacteria showed that contrary to assumptions that switching events are independent and Poisson distributed, there is a CW bias due to inherent stochastic fluctuations in the signalling network. This CW bias manifests as deviations from the expected flat power spectrum and a cumulative power law distribution of CCW intervals [16]. Building on this experimental input of CW bias due to fluctuations in the signalling pathway, simulation studies predicted a power law dependence of bacterial runlengths even in the absence of any chemoattractant. This suggests that a Levy-walk strategy might be more beneficial for bacteria in such fluctuating environments [17]. Recent experimental analysis of single bacterial trajectories in 3D homogeneous environments suggests that run length distributions are indeed power-law distributed, and the trajectories are thus superdiffusive, even in absence of chemoattractant [18]. However, the bacterial motion in biological contexts in often characterised by confinement, and also presence of chemical gradients, and it remains an open question as to how bacteria modulate their swimming strategies in such environments.

We have developed a novel two-layer microfluidic device to study the combined effect of the degree of confinement and a chemical gradient on the motility of *E. coli*. Our device also allows us to study this effect in two dimensions, in two different channel widths simultaneously, while maintaining a uniform fluidic resistance. This device has significant utility in understanding the evolution of bacterial cell motility in response to different confined conditions and chemical cues. By tracking the motion of individual *E. coli* in 50 *μ*m and 10 *μ*m wide channels, in the presence and absence of chemical gradients, we quantify how *E. coli* adapts its swimming strategies to these differing physical and chemical cues. We show that bacterial trajectories in such channels are superdiffusive, with power law distributed runlengths, even in absence of chemical gradients. Introduction of a chemical gradient, or stronger confinement increases the exponent of the superdiffusive behaviour, and decreases the power law exponent of runlengths, with increasing probability of longer runs. Further, both confinement and chemoattraction suppress large angle tumbles, indicating bacteria adopt both these strategies of modulating runs and tumbles to adapt their swimming strategies.

## I. EXPERIMENTAL METHODS

### A. Equipment and chemicals

2-inch diameter silicon wafers were purchased from Prolyx Microelectronics (Bengaluru, India). SU-8 2002 and SU-8 2025 were obtained from Microchem Corporation (Westborough, USA). A Karl Suss MJB4 mask aligner was used for the fabrication of SU-8 masters on silicon substrates by lithography. The height of the features was characterized using an Ambios-XP2 profilometer (Ambios Technology Inc., USA). Sylgard 184 elastomer (polydimethylsiloxane, PDMS) kit was obtained from Dow Corning (Michigan, USA). Inlets and outlets were punched in the PDMS devices using a 1.5 mm diameter biopsy punch (Med Morphosis LLP, India). The PDMS devices were bonded to 22 mm diameter circular (no. 1.5) glass cover slips (Blue Star) using a PDC-32G-2 cleaner from Harrick Plasma, USA. An Eclipse Ti-U series inverted fluorescence microscope (Nikon corporation, Japan) was used to capture the images of the FITC-dextran gradient profiles in our device. A spinning disk confocal microscope (Carl Zeiss) was used for capturing the time lapse images of bacteria inside the microfluidic device.

LB broth (catalog no. M575-500G), Phosphate buffered saline (PBS) (catalog no.TS1006), Bovine Serum Albumin (BSA) (catalog no. MB083-100G) and glucose (catalog no. RM10215-25G) were purchased from Himedia, India. 70 KDa FITC-dextran used to characterize the gradient profile was purchased from Sigma-Aldrich (catalog no. 46945, India).

### B. Design of the microfluidic device

As shown in Fig. 1A and 1B, the device design resembles a ladder. The side-rails of this ladder consist of two parallel feeding channels (inducated in green) of 100 *μ*m width and 25 *μ*m height. Each feeding channel is connected to an inlet and an outlet. The two feeding channels are connected to each other by sixteen parallel microchannels (indicated in blue) of 2 *μ*m height, forming the rungs of the ladder. These rungs are henceforth called ’motility observation channels’ or MOCs. Eight of the MOCs are 50 *μ*m wide, while the other eight channels are 10 *μ*m wide. The 50 *μ*m MOCs have a length of 2095 *μ*m, while the 10 *μ*m MOCs are 376 *μ*m long. The different channel lengths ensure that the hydraulic resistances (*R_h_*) of the 50 *μ*m and the 10 *μ*m MOCs are equal. This design minimizes any bacterial motility due to fluid flow and allows us to measure the active motility of bacteria.

**FIG. 1:**
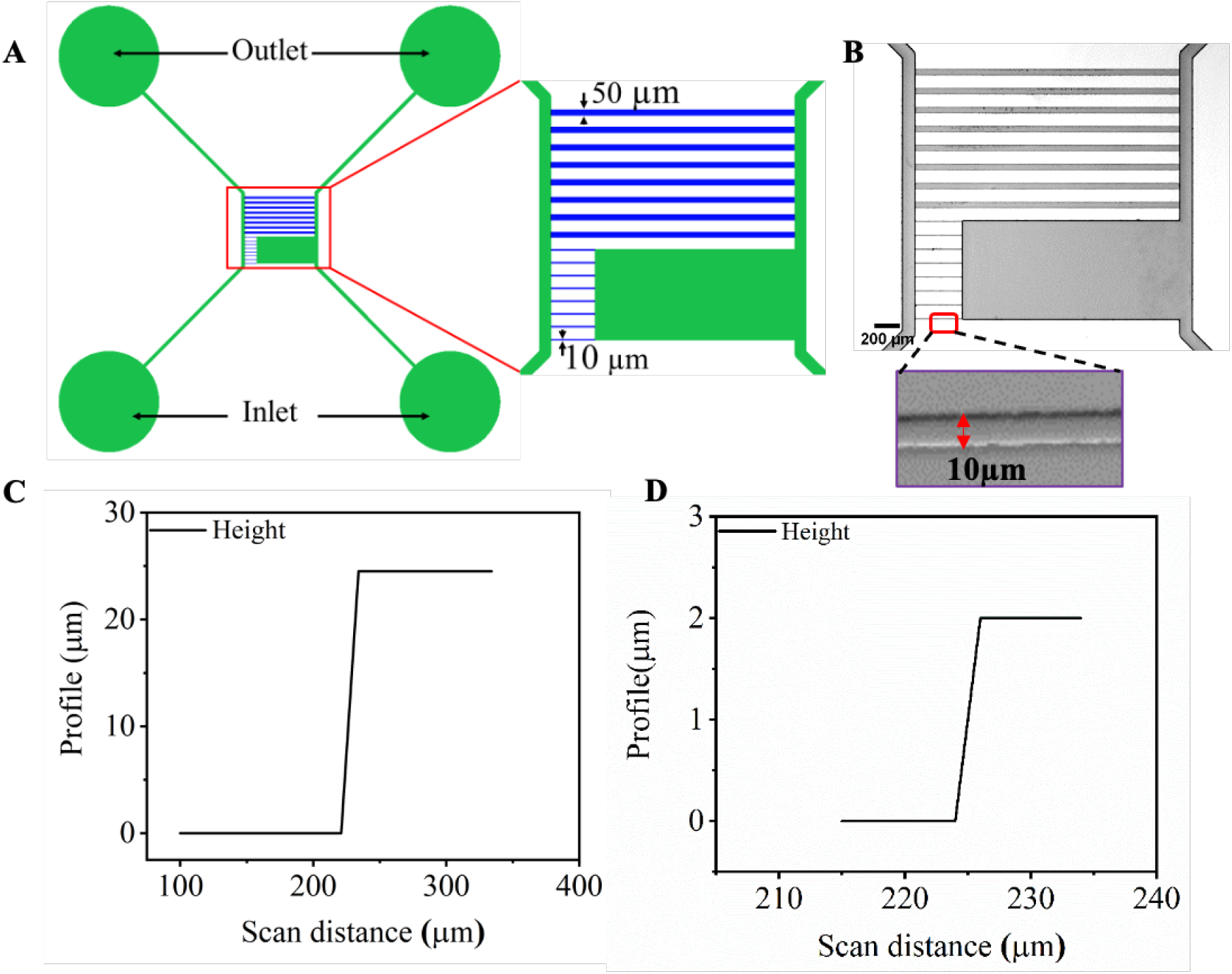
(A) SChematic diagram of the microfluidic device. Inlets, outlets and feeding channels are indicated in green. 50 *μ*m and 10 *μ*m wide motility observation channels (MOC) are indicated in blue. (B) Microscopic image of the device, with the inset showing a 10 *μ*m wide MOC. (C, D) Profilometry data showing the heights of the feeding channel and the MOC respectively.

### C. Fabrication of the microfluidic device

The mask was designed in CleWin (version 5). The chrome-on-glass mask was fabricated by laser writing at the nanofabrication facility of Indian Institute of Science, Bengaluru. The master was patterned by two-step lithography using SU-8 2002 (for MOCs) and SU-8 2025 (for feeding channels) on a silicon substrate. Initially, SU-8 2002 was spin-coated on a silicon wafer with a spreading spin of 500 rpm for 5 sec (with 100 rpm/sec acceleration) and a final spin of 2000 rpm for 30 sec (with a 300 rpm/sec acceleration). The wafer was soft-baked at 95 °C for 3 min and exposed under UV for 1.8 sec. The exposed wafer was baked at 95 °C for 5 min and developed using the proprietary SU-8 developer. After rinsing and drying, the height of the feature was measured using profilometry. Next, the feeding channels were patterned using SU-8 2025 with the same spin coating parameters. The coated wafer was soft-baked at 95 °C for 5 min, exposed under UV for 6 sec and post-baked at 95 °C for 5 min. Following development, rinsing and drying, profilometry was performed to check the height of the feeding channels. The microfluidic devices were made using PDMS following standard soft lithography protocols. Inlets and outlets were punched using a 1.5 mm diameter biopsy punch. The devices were finally plasma-bonded to clean glass coverslips.

### D. Characterization of the gradient profile using FITC-dextran

We characterized the gradient profile inside the MOCs using 70 KDa FITC-dextran. The device was filled with fresh Luria-Bertoni (LB) broth. We then pipetted 10 *μ*l of 1 mM FITC-dextran (prepared in 1X PBS) into the right-hand feeding channel. Time lapse images of diffusing FITC-dextran in the MOCs were captured at 1 fps using a 4X objective.

We quantified the gradient profiles at different time intervals up to 560 sec to understand when the gradient becomes stable in each channel. Since there is a single source of FITC-dextran in the right feeding channel, we expect the intensity profile along any MOC to decay exponentially. We found that the intensity is saturated in the feeding channels. Therefore, we defined the source (*x*_0_) at a distance of 50 *μ*m and 125 *μ*m from the feeding channel in the 50 *μ*m wide and the 10 *μ*m wide MOC, respectively. Intensity values along the MOCs were normalized by dividing them by the intensity value at the source (i.e. at *x*_0_). We plotted the normalized intensity profiles at different time points, as shown in Fig. 2A-B. The intensity profiles were fitted with the equation 1.

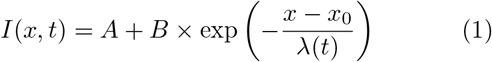

where *A* is the background intensity, *B* is the source intensity, *x*_0_ is the distance of the source from the feeding channel and λ is the characteristic length scale.

**FIG. 2:**
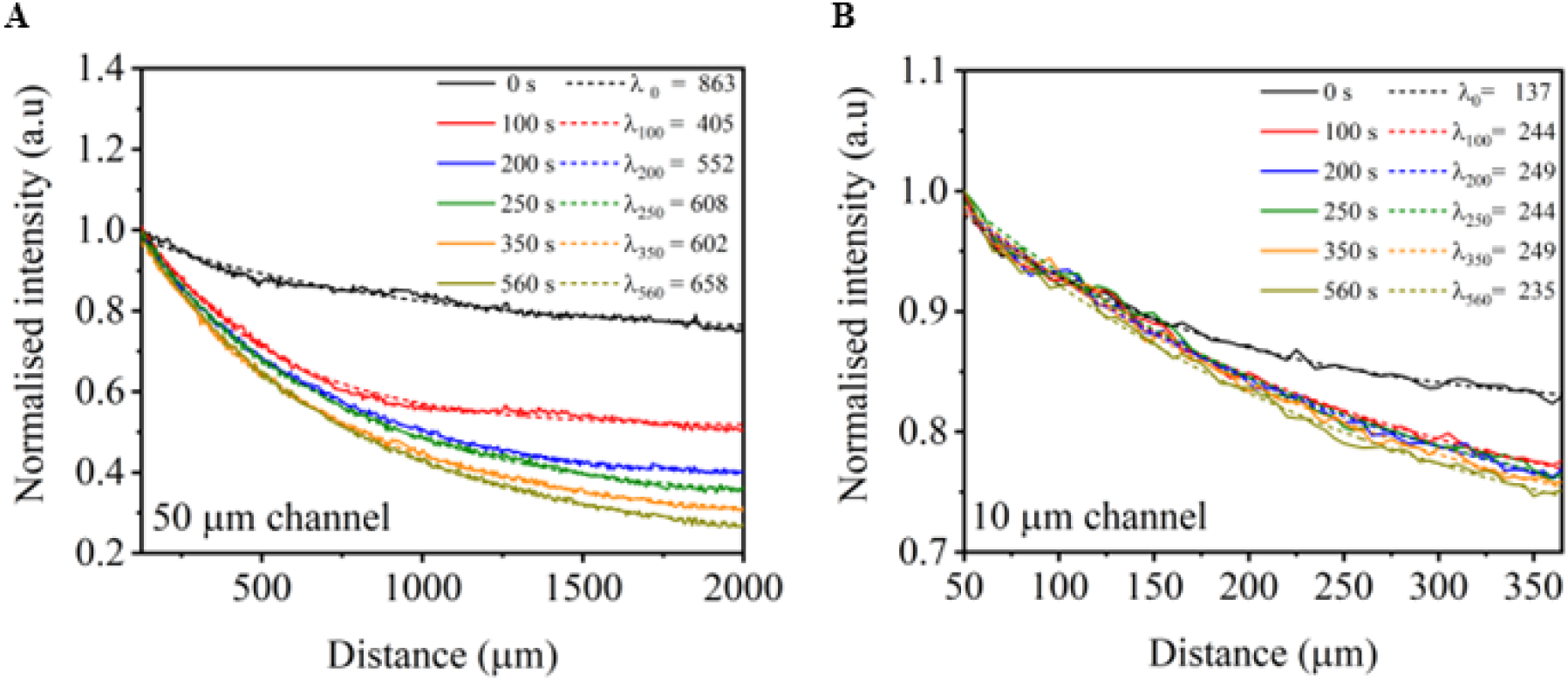
Gradient profiles of FITC-Dextran in (A) 50 *μ*m and (B) 10 *μ*m MOCs. Each solid curve represents the average gradient profiles from four MOCs. The dotted lines show the exponential fit, with λ indicating the characteristic length scale.

The extracted characteristic length scales λ(*t*) are shown for five representative time points as shown in the figure 2A-B. The steady state values of λ (i.e. λ_*ss*_^50^) for the 50 *μ*m channel is 605 *μ*m, while λ_*ss*_^10^ for the 10 *μ*m channel is 244 *μ*m. We observed a stable gradient after 200 sec in the 50 *μ*m channel, whereas a stable gradient was established after 100 sec in the 10 *μ*m channel.

### E. Passivation of the microfluidic device

We passivated the channels to prevent adhesion of *E. coli* to PDMS or glass during motility experiments. We filled the devices with a 10 mg/mL solution of bovine serum albumin (BSA) in distilled water. The devices were incubated with this solution at 37°C for at least one hour.

### F. Bacterial culture

The bacterial strain CGSC 8237 (MG1655 wild type K12 motile strain, isolated from a mixed population of CGSC 6300) was purchased from the Yale University’s Coli Genetic Stock Center. This strain of *E. coli* harbors a vector plasmid carrying nptII and GFP markers. We cultured the *E. coli* in LB broth supplemented with 40 *μ*g/ml kanamycin. We first cultured the bacteria overnight at 37°C, with 180 - 200 rpm agitation. We then sub-cultured it in 1 ml fresh LB broth until it reached mid-log phase (0.4× 10^9^ cfu/ml). The suspension was centrifuged at 5000 rpm for 10 min to get a bacterial pellet. This pellet was washed three times in 1X PBS and re-suspended in fresh LB broth. The suspension was diluted 10 to 200 times with LB broth for the motility experiments.

### G. Motility experiments

The microfluidic chip was mounted on the thermal stage of a spinning disc confocal microscope held at 37° C. The entire chip was filled with fresh LB broth. We then pipetted 10 *μ*l bacterial suspension into the inlet of the left feeding channel while removing the excess liquid from the corresponding outlet. All four inlets and outlets were then sealed with Scotch tape to prevent fluid evaporation. Bacteria swam into the motility observation channels where they were imaged. For experiments with the chemoattractant, we added 10 *μ*l of 1 mM glucose into the inlet of the right feeding channel before adding bacteria into the left feeding channel. All motility measurements were performed under static flow conditions.

### H. Time lapse microscopy and image analysis

Time lapse images of bacteria were captured at 50 ms intervals using a 40X (1.5 NA) oil immersion objective of the spinning-disk confocal microscope. The captured images were analyzed using MATLAB (R2020a) and ImageJ.

## II. RESULTS AND DISCUSSION

### A. Both confinement and chemoattractant reduces wall crawling of *E. Coli*

Bacterial motility in less confined spaces is predominantly controlled by hydrodynamic forces. But in case of mesoscale-sized channels, it is also influenced by wall interactions [14]. Tokarova et al.(2021) and Vizsnyiczai et al.(2020) have demonstrated that *E. coli* prefer swimming along the walls in wider channels. But they swim axially in narrow (*w* < 2.5 – 6 *μ*m) microchannels due to steric hindrance[14][15].

We use a MATLAB code to extract the XY-coordinates of bacterial cell centroids across our microchannels at regular time intervals. We plot these coordinates as a function of time to visualize the trajectories of the *E. coli* in our channels. As shown in Fig. 3A and 3C, the trajectories indicate that the *E. coli* prefers swimming along the walls in both 50 *μ*m and 10 *μ*m channels. But the presence of glucose reduces this wall-swimming behavior in both channels (Fig. 3B and 3D). The shift of cell movement towards the channel center is more pronounced in the 10 *μ*m channel.

**FIG. 3:**
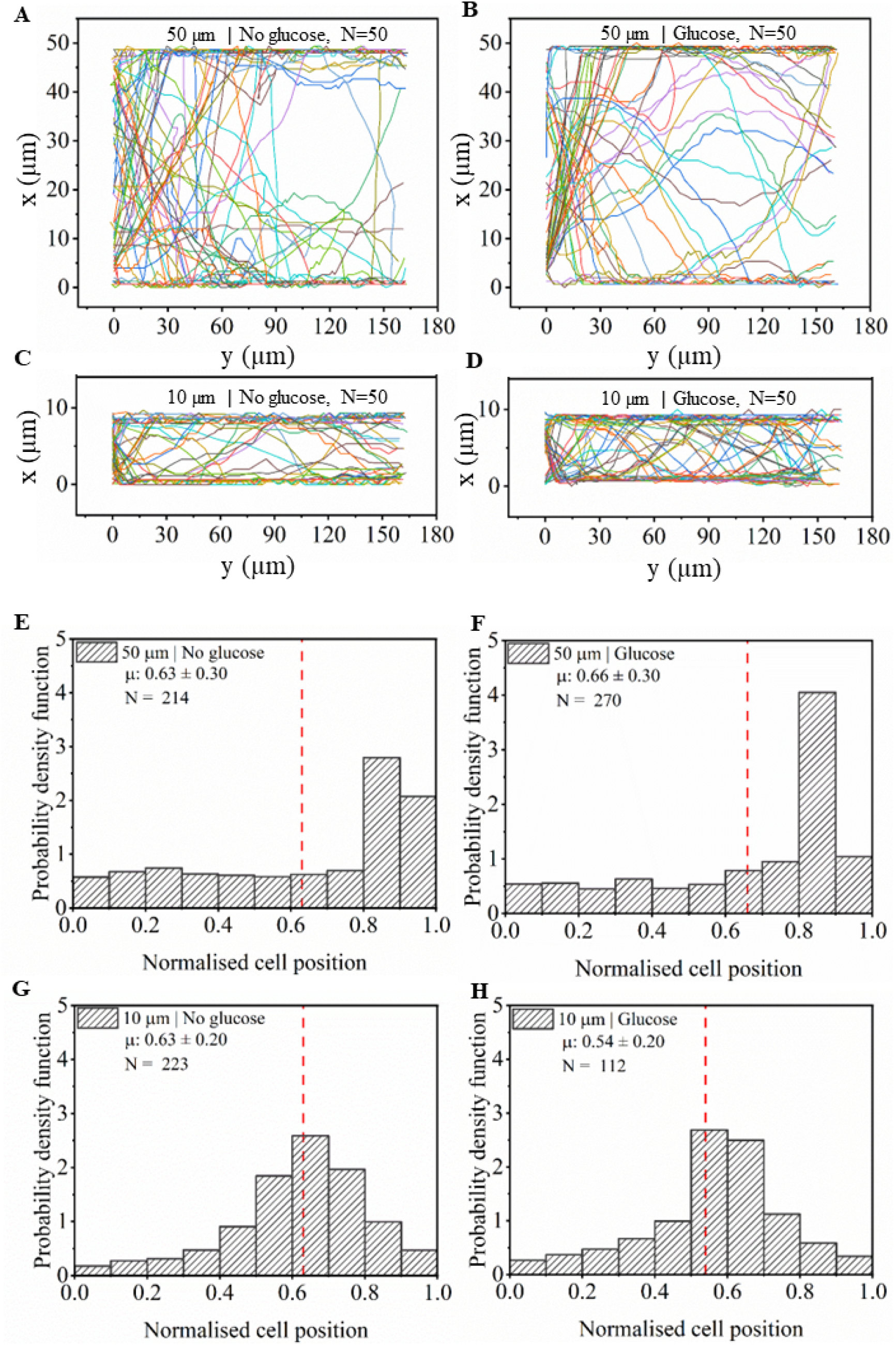
(A,B,C and D) Representative trajectory in 50 *μ*m and 10 *μ*m channels. (E,F,G and H) Normalised cell position in 50 *μ*m and 10 *μ*m channels. Vertical red dots = mean ± SD, N = No. of trajectory.

To quantify this shift in motility patterns, we obtain the positions of the cell centroids of each bacteria for the entire duration of its trajectory. We then plot the probability distribution of these cell positions by normalising with the channel width (*w* = 50 *μ*m and 10 *μ*m) to infer the locations where bacteria spend maximum time along the channel width. A normalised cell position of zero indicates pure axial swimming, while one indicates complete wall crawling on either boundary wall. As shown in Fig. 3E, most of the *E. coli* in the 50 *μ*m channels favor positions close to the walls in the absence of chemical cues. In the presence of glucose, there is no significant change in the mean position (Fig. 3G). However, as shown in figure 3B, there is a reduction in the number of bacterial trajectories which are maximally close to the wall in the presence of glucose. In case of the 10 *μ*m channels, the *E. coli* swim further away from the walls due to confinement (Fig. 3C). On addition of glucose, the distribution as well the mean position shifts further towards the center, as shown in Fig. 3H.

The effect of confinement on cell motility has been previously reported[14][15][19]. In particular, strong confinement (*w* < 2.5 – 4 *μ*m) has been shown to shift the bacterial populations from a wall crawling to an axial swimming behaviour in both motile and non-tumbling *E. coli* [14][15]. This is consistent with our observation that in comparison to trajectories in a 50 *μm* channel, bacteria are found more to the center in 10 *μm* channels. However, the combined effect of confinement and chemoattractant has not been reported earlier. We show that for moderate confinements, chemoattractant induces a shift in the average cell position towards the center of the channel, and opens the possibility that transitions from wall crawling to axial swimming can be induced by strong chemoattractant gradients.

### B. Bacterial motion is super-diffusive under confinement and chemoattraction

To characterise the transport coefficients, we analyse the mean square displacement (MSD) of the *E. coli* within our different channels geometries. The MSD is fitted to a power-law form [20][21],

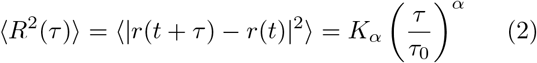

where *τ*_0_ is some reference time. For normal diffusive transport, the power-law exponent *α* = 1, and the prefactor *K*_*α*=1_ then determines the standard diffusion coefficient, *K*_*α*=1_ = 2*dτ*_0_*D* in *d*–dimensions [22][23]. For anomalous transport, where *α* ≠ 1, the use of the dimensionless time parameter *τ*/*τ*_0_ allows us to define an effective generalised diffusion coefficient *D_α_* = *K_α_*/*τ*_0_ [22][23]. Throughout this manuscript, we use a reference timescale of *τ*_0_ = 1s to report effective diffusion coefficients across different confinements and chemical gradients with different values of the anomalous diffusion exponent *α*.

We first characterise the transport for *E. coli* trajectories in the bulk feeding channels (Fig. 4a). Contrary to expectations, and in line with recent reports [18], the trajectories were super-diffusive at all timescales observed in our exponenents. At long times, we fit the MSD to Eq. 2 to obtain a superdiffusive exponent of *α* = 1.47 (Fig. 3A). The effective diffusion coefficient in this case is *D_α_* = 331.1*μm*^2^/*s*.

**FIG. 4:**
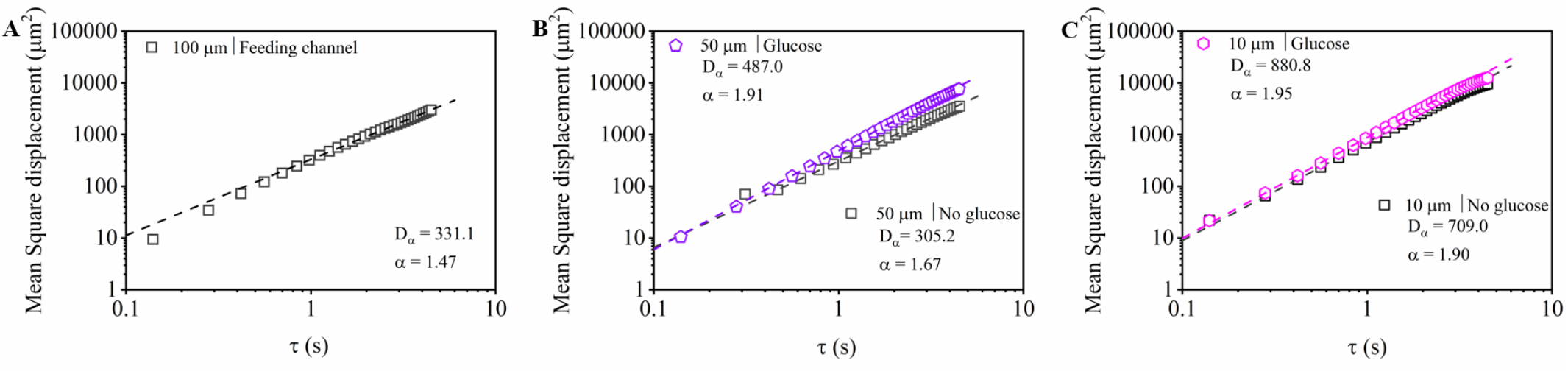
Mean square displacement in (A) Bulk feeding channel (B) 50 *μ*m channel and (C) 10 *μ*m channel. *D_α_* (*μm*^2^/*s*) and *α* are the generalised diffusion constant and power law exponent respectively. Dotted lines indicate fits to the experimentally obtained data points.

Next we analyse the trajectories in the 50 *μ*m channel (Fig. 4B). In the absence of chemoattractant, simple confinement induces deviations from the behaviour observed in the bulk feeding channels, with an exponent *α* = 1.67 indicating stronger superdiffusive transport even with this weak confinement. Previous studies on bacterial motion in pore (spherical) confinement showed an initial super-diffusive motion transitioning to a sub-diffusive motion for long lag times (*τ* > 1 – 10s) [13]. In contrast, for our channel geometries, for the range of lag times (*τ* ~ 10s) investigated, the motion is consistently super-diffusive. On introducing a chemoattractant, the motion of *E. coli* approaches ballistic transport, with *α* = 1.91.

A previous study by Yang et al.(2015)[24] shows that *E. coli* cells exhibit a ballistic to diffusive transition in the presence of chemical cues, similar to transport in the absence of chemoattractant. The initial ballistic or superdiffusive regime corresponds to single runs of bacteria, and in the presence of chemoattractant, this regime persists for longer time. In our experiments, in the presence of chemoattractant, we observe superdiffusive motion for all time intervals *τ* ~ 10s that were investigated. Curiously, the mean duration of individual runs did not change appreciably on addition of chemoattractant (Suppl. Fig. S1) even though the transport approaches the ballistic regime.

We next study the MSD of trajectories in stronger confinement (10 *μ*m channels, both with and without chemoattractant (Fig. 4C). For these values of confinement, even in the absence of chemoattractant, the motion is near-ballistic, with an exponent of *α* = 1.90 which increases slightly to *α* = 1.95 on introduction of glucose. In this case, mean run times are also much longer, ~ 2.8 – 3.3sec, (Suppl. Fig. S1), and thus the MSD plots capture the run phase of individual *E. coli* in the regimes of lag time investigated. This was also consistent with our observation of *E. coli* trajectories in 10 *μ*m channels, where the channel length was often traversed without any large angle tumbles. Note that while the exponent *α* did not vary appreciably on addition of chemoattractant, the effective diffusion coefficient *D_α_* increased on addition of glucose, indicating higher average speeds.

### C. Both confinement and chemoattractant increase cell speeds and probability of long runs

Previous studies have reported that confinement and chemical cues influence cell speeds and run lengths[7][19][12]. In our study, we used the XY-coordinates of *E. coli* (as obtained using our MATLAB code) to quantify cell speeds and run lengths. We first identified individual runs from bacterial trajectories by quantifying the angular deviation between cell positions at three successive time instants. If these deviations were less than a threshold of ±20° they were considered to be part of the same run. If the angle of motion is greater than the threshold, we categorized it as a “tumble”. By this method, we identified individual run lengths in different experimental configurations. We divided these individual runlengths by the duration of each run to obtain cell speeds in the run-phases.

The distributions of cell speeds are shown in Fig. 5A–D. Increasing the confinement increases the mean speed from 16.5 ± 7.4*μ*m/sec in the 50 *μ*m channels to 26.5 ± 7.4*μ*m/sec in the 10 *μ*m channels. The mean speeds also increase on addition of chemoattractant, as has been reported in the literature [25], with a 1.4-fold increase for 50 *μ*m channels, and a 1.1-fold increase for 10 *μ*m channels.

**FIG. 5:**
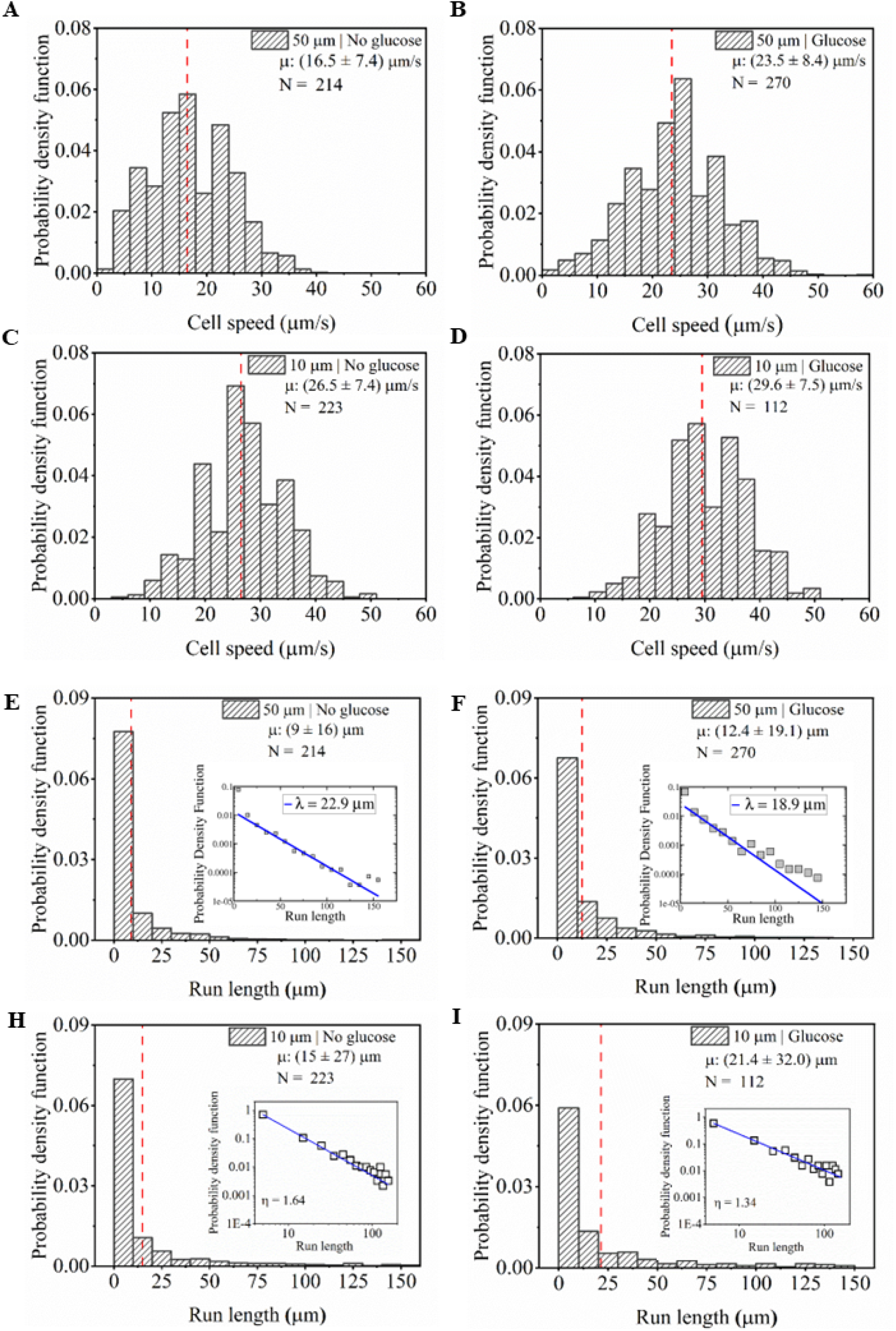
(A,B,C and D) Speed distributions in 50 *μ*m and 10 *μ*m wide channels in absence (A and C) and presence (B and D) of chemoattractant. (E,F,G and H) Run lengths in 50 *μ*m and 10 *μ*m wide channels in absence (E and G) and presence (F and I) of chemoattractant. Dotted red lines indicate the mean ± SD. N indicates the number of trajectories in each experiment. The insets show the fit (blue lines) to the run length distributions. The data for 50 *μ*m wide channels show an exponential fit, while the data in the 10 *μ*m channels show a power law fit.

Figures 5E – H show the effects of confinement and glucose on the typical run lengths of *E. coli*. The distributions of runlengths for both confinements and in the presence and absence of chemoattractant are shown in Fig. 5E-H. We found that the mean run length increases with increase in confinement, as well as with the introduction of glucose. In line with classical expectations[7], we find the distribution of runlengths for weak confinement (50*μm* channels) in the absence of glucose is exponential (Fig. 5E inset), with a characteristic runlength of 22.9*μm*. On introduction of a glucose gradient in this channel, strikingly, the run length distribution deviates from a pure exponential, with an enhanced probability for longer runs (Fig. 5F inset). This deviation from exponential behaviour is in line with observations of nonexponential distributions of intervals of CW and CCW rotations in immobilised *E. Coli* [16], and with recent reports of non-exponential runlength distributions in 3D solutions [18]. Having established deviations from exponential behaviour in presence of chemoattractant gradient, we now turn to the effect of stronger confinement on runlength distributions. For a 10*μm* confinement, even in the absence of chemoattractant, the effect is even more drastic. The exponential distribution of runlengths observed in 50*μm* channels shifts to a power law distribution in this case (Fig. 5G inset). We fit a power law of the form ~ *x^−η^* to the runlength distribution, with a fitted exponent *η* = 1.64 in this case. Power-law distributions in bacterial runlengths have been hypothesized to occur due to fluctuations in the signalling pathway, and we demonstrate for the first time that confinement can also induce deviations from ideal swimming behaviour, with a swimming strategy that favours longer runs. As expected, on introduction of a gradient in glucose concentration in the 10 *μ*m channels, the power law behaviour persists, however longer runs become even more likely, with a consequent further decrease in the power law exponent, *η* = 1.34 in this case (Fig. 5H inset).

### D. Confinement and chemoattractant decrease E. coli tumbles, and specifically suppress large angle tumbles

We used tumbling frequency to understand the effect of confinement and chemoattractant on the directional angle of *E. coli* motility [25][12]. As discussed earlier, tumbles were defined when the change in angular direction between three successive time points was greater than 20°. Very small threshold angles overestimate tumbles, while larger threshold angles may miss transitions between different run phases (see Suppl. Fig. S2). For each individual trajectory of *E. coli*, we identified the number of tumbles in the duration the bacteria remained in the field of view. Thus, we identified a tumbling frequency from each trajectory by dividing the number of tumbles by the duration. Data from multiple such trajectories were then used to generate a distribution of tumbling frequencies.

As shown in figures 6A – D, increasing the confinement from 50 *μ*m to 10 *μ*m reduces the tumbling frequency of *E. coli*, with many bacteria traversing the 10 *μ*m channel without any apparent tumbles. The introduction of glucose in both channels also decreases the tumbling frequency.

**FIG. 6:**
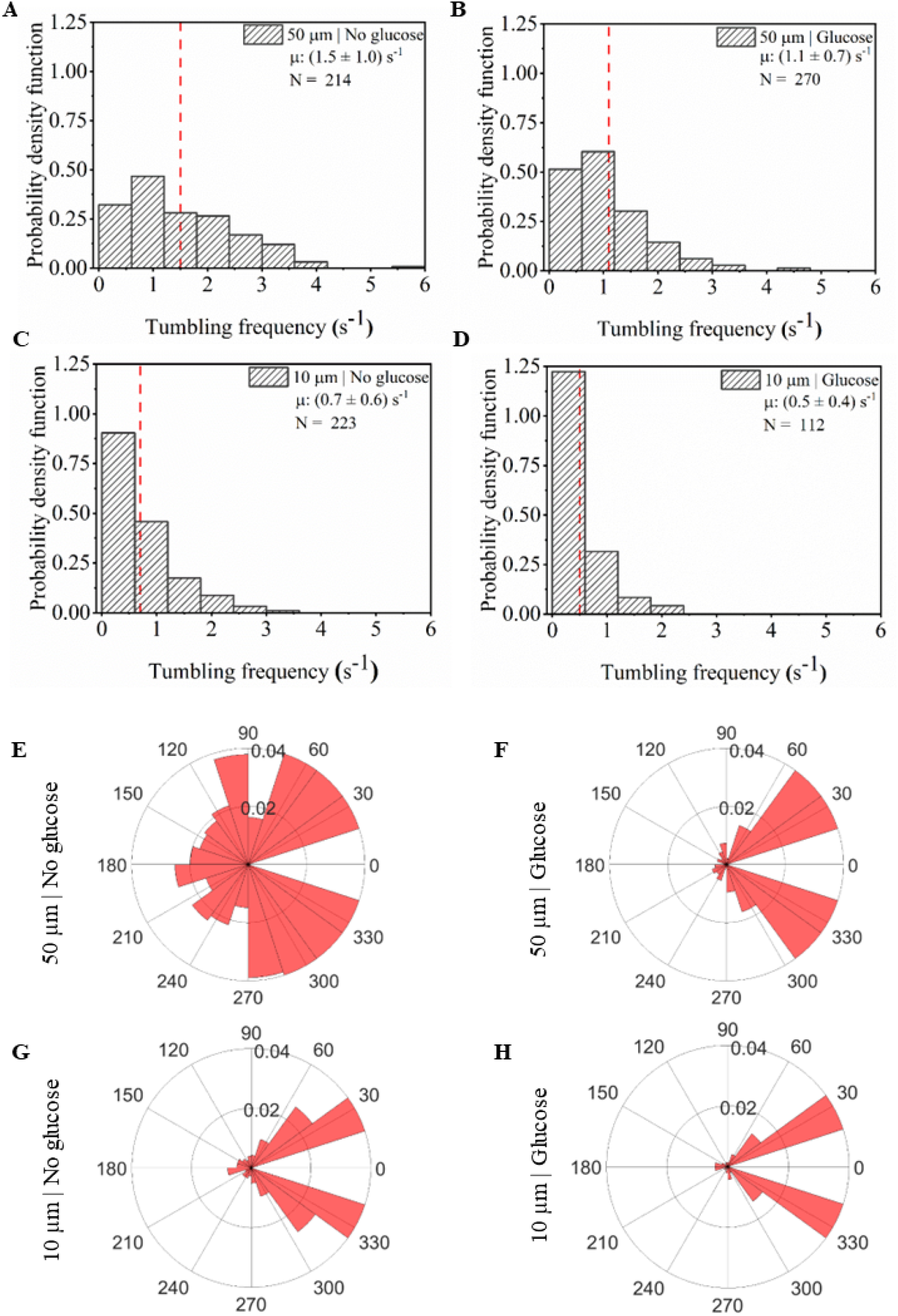
(A,B,C and D)Tumbling frequency (E,F,G and H) Rose plot for turn angle distribution in 50 *μ*m and 10 *μ*m width channel. Vertical red dots = mean ± SD, N = No. of trajectory.

Evidence in the literature suggests that chemoattractant not only increases typical run durations, but also changes the distribution of tumbling angles, with a much reduced probability for large angle tumbles in the presence of a chemical cue. This is consistent with our observations, both for the 50 *μ*m and the 10 *μ*m channels. As can be seen in Fig. 6F and H, most tumbles in the presence of glucose are in a ±60° window, compared to the large angle tumbles in Fig. 6E. Interestingly, confinement produces a similar effect on tumbling angles, suppressing large angle tumbles in the 10 *μ*m channels, as can be seen in Fig. 6G.

## III. CONCLUSION

In summary, we compared how *E. coli* adapts its motility in response to confinement and chemical gradient. We designed a two-layer microfluidic device to study these effects under quasi-2D conditions in absence of any drift due to fluid flow. We found that the trajectories of *E. coli* are superdiffusive. Both confinement and chemical gradient increased the exponent of the superdiffusive behaviour with a higher probability of longer runs. For weak confinement in absence of gradient, runlengths are exponentially distributed. Both confinement and chemical gradient induce deviations from this exponential behaviour, with the runlengths following a power law behaviour. We also noted a suppression of large angle tumbles in presence of either confinement or a chemoattractant. Our observations indicate that *E. coli* modulates both its runs and tumbles inside microfluidic confinement or in presence of a chemical gradient. While changes in the signalling pathway can be related to the motility of *E. coli* in response to chemical cues, similar studies on the response of *E. coli* to physical cues (confinement) can provide interesting insights into how bacteria navigate their natural habitats.

## Data and code availability

MATLAB code and experimental datasets are available on request. Please contact the corresponding author.

## ACKNOWLEDGEMENTS

MRR acknowledges a PhD fellowship (DBT/2017/IITB/959) from the Department of Biotechnology, Government of India. JEG acknowledges a postdoctoral fellowship from the Indian Institute of Technology Bombay. The authors acknowledge the IIT Bombay’s Nanofabrication Facility and the Spinning Disc Confocal Microscopy Facility of IIT Bombay. The authors also thank Pradip Shinde for help with confocal imaging, and Dr Shilpi Pandey and Santosh Jinnawar for other technical and administrative assistance.

## Author contributions

MRR, DP and MKM designed the research plan. MRR performed the experiments. MKM, JEG and SK wrote the MATLAB code for image analysis. MRR, JEG and MKM performed the data analysis. MRR, MKM and DP wrote the manuscript.

## Supplementary Information

**Figure S1:**
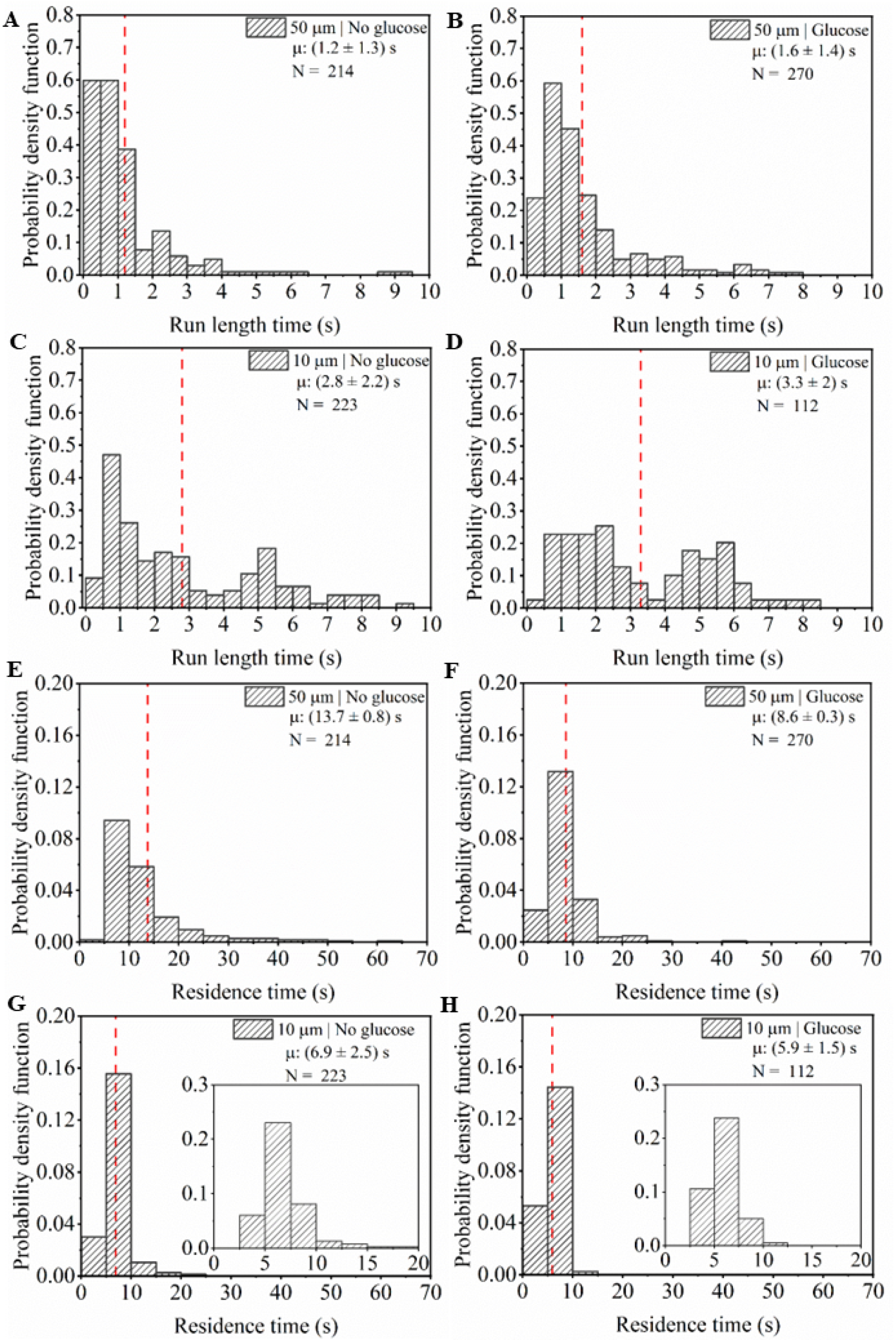
Distributions of (A,B,C and D) Run length time, and (E,F,G and H) Residence time in 50 *μ*m and 10 *μ*m width channel. Vertical red dashed line = mean ± SD, N = No. of trajectory.

**Figure S2:**
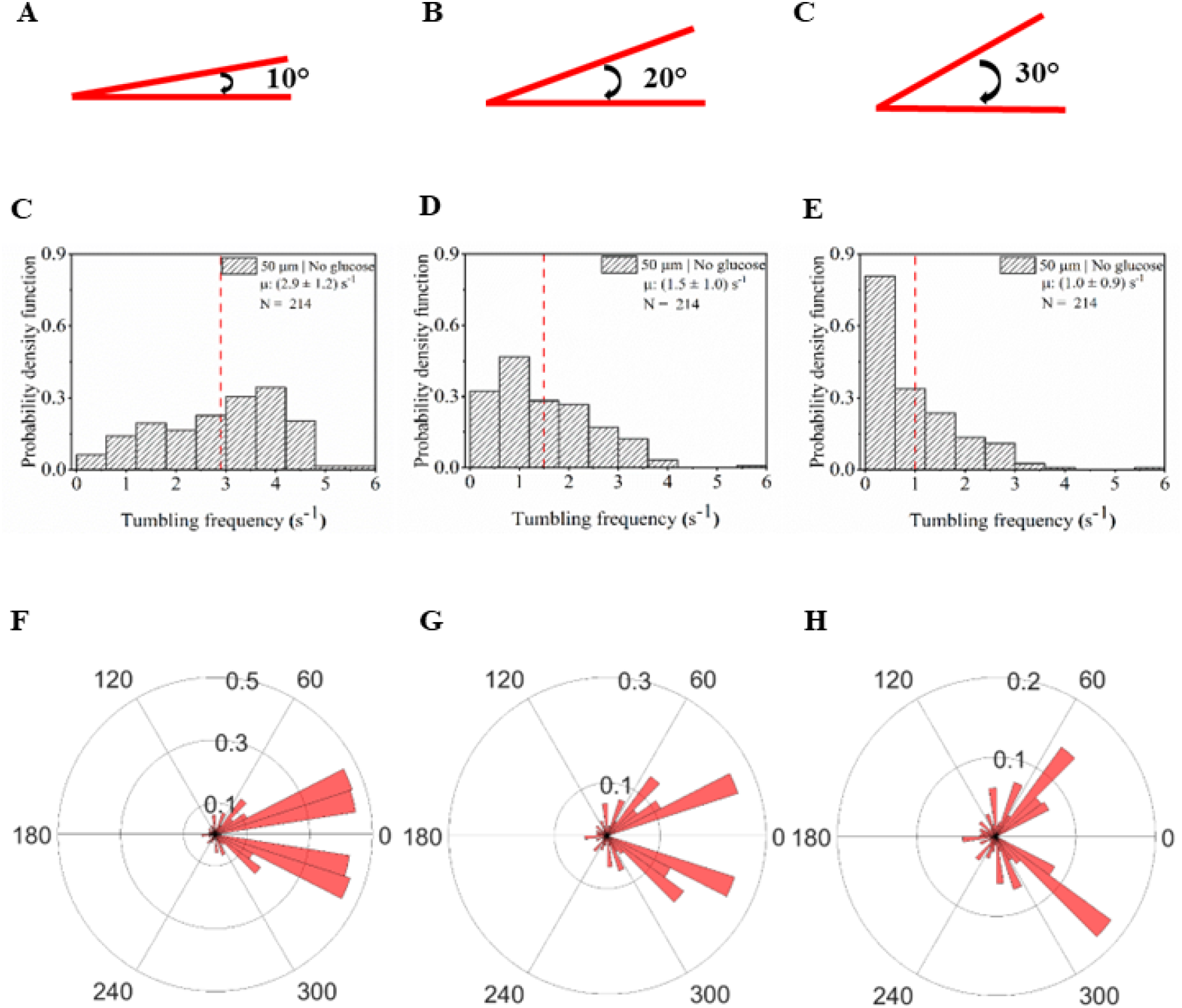
(A-H) Sensitivity of tumbling frequency distribution and Rose plot of tumble angles on choice of threshold angle for defining tumbles.

